# CAMSA: a Tool for Comparative Analysis and Merging of Scaffold Assemblies

**DOI:** 10.1101/069153

**Authors:** Sergey S. Aganezov, Max A. Alekseyev

## Abstract

**Motivation:** Despite the recent progress in genome sequencing and assembly, many of the currently available assembled genomes come in a draft form. Such draft genomes consist of a large number of genomic fragments (*scaffolds*), whose positions and orientations along the genome are unknown. While there exists a number of methods for reconstruction of the genome from its scaffolds, utilizing various computational and wet-lab techniques, they often can produce only partial error-prone scaffold assemblies. It therefore becomes important to compare and merge scaffold assemblies produced by different methods, thus combining their advantages and highlighting present conflicts for further investigation. These tasks may be labor intensive if performed manually.

**Results:** We present CAMSA—a tool for comparative analysis and merging of two or more given scaffold assemblies. The tool (i) creates an extensive report with several comparative quality metrics; (ii) constructs the most confident merged scaffold assembly; and (iii) provides an interactive framework for a visual comparative analysis of the given assemblies. Among the CAMSA features, only scaffold merging can be evaluated in comparison to existing methods. Namely, it resembles the functionality of assembly reconciliation tools, although their primary targets are somewhat different. Our evaluations show that CAMSA produces merged assemblies of comparable or better quality than existing assembly reconciliation tools while being the fastest in terms of the total running time.

**Availability:** CAMSA is distributed under the MIT license and is available at http://cblab.org/camsa/.

## 1. Introduction

While genome sequencing technologies are constantly evolving, researchers are still unable to read complete genomic sequences at once from organisms of interest. So, genome reading is usually done in multiple steps, which involve both *in vitro* and *in silico* methods. It starts with reading small genomic fragments, called *reads*, originating from unknown locations in the genome. Modern shotgun sequencing technologies can easily produce millions of reads. The problem then becomes to assemble them into the complete genome. Existing de novo genome assembly algorithms can usually assemble reads into longer genomic fragments, called *contigs*, that are typically interweaved in the genome with highly polymorphic and/or repetitive regions. The next step is to construct *scaffolds*, i.e., sequences of (oriented) contigs along the genome interspaced with gaps. The last but not least step is genome finishing that recovers genomic sequences inside the gaps within the scaffolds.

Unfortunately, the quality of scaffolds (e.g., exposing severe fragmentation) for many genomes makes the finishing step infeasible. As a result, the majority of currently available genomes come in a *draft* form represented by a large number of scaffolds rather than complete chromosomes [35]. This emphasizes the need for improving the assembly quality of genomes by constructing longer scaffolds from the given ones,^1^ which we refer to as the *scaffold assembly problem*. In other words, the scaffold assembly problem asks for reconstruction of the order of input scaffolds along the genome chromosomes.

A number methods have been recently proposed to address the scaffold assembly problem by utilizing various types of additional information and/or *in vitro* experiments. These methods are based on jumping libraries [20, 40, 22, 17, 27, 11, 7], long error-prone reads (such PacBio or MinION reads) [45, 5, 6, 8, 24], homology relationship between multiple genomes [3, 1, 2], wet-lab experiments such as the fluorescence *in situ* hybridization (FISH) [36, 41], genome maps [32, 42, 28], higher order chromatin interactions [9], and so on. Depending on the nature and accuracy of utilized information and techniques, assemblies produces by different methods may still be incomplete and contain errors, thus deviating from each other. Moreover, some scaffolding methods (e.g., based on FISH or HiC data) can produce assemblies, where the (strand-based) orientation of some assembled scaffolds is yet to be determined.

It therefore becomes crucial to determine what parts of different assemblies are consistent with and/or complement each other, and what parts are conflicting with other assemblies (or even within the same assembly). Furthermore, some scaffold assemblies may utilize only a fraction of the input scaffolds (e.g., homology-based assembly methods do not take into account unannotated scaffolds), thus posing a problem of analyzing and comparing assemblies of varying subsets of scaffolds. Comparative analysis of scaffold assemblies produced by different methods can help the researchers to combine their advantages, and highlight potential conflicts for further investigation. These tasks may be labor-intensive if performed manually.

While there exists a number of methods [47, 48, 34, 43, 30, 46, 44] for reconciling multiple assemblies of the same organism, they all are limited only to oriented scaffolds and thus are inapplicable to scaffold assemblies that include unoriented scaffolds. Furthermore, some of these methods require a reference genome sequence, which is often unavailable for non-model organisms. On the other hand, reconciliation methods that operate in de-novo fashion often process the input assemblies progressively, which makes such methods sensitive to the order of the input assemblies and affects the quality of the reconciled assembly.

We present CAMSA, a tool for comparative analysis and de-novo merging of scaffold assemblies. CAMSA takes as an input two or more assemblies of the same set of scaffolds and generates a comprehensive comparative report for them. Input assemblies can include both oriented and unoriented scaffolds, which enables CAMSA to process assemblies from the full range of scaffolding techniques (both *in silico* and *in vitro*). The generated comparative report not only contains multiple numerical characteristics for the input assemblies, but also provides an interactive framework, allowing one to visually analyze and compare the input scaffold assemblies at regions of interest. CAMSA also computes a *merged assembly*, combining the input assemblies into a more comprehensive one that resolves conflicts and determines orientation of unoriented scaffolds in the most confident way. The non-progressive nature of merging in CAMSA eliminates the dependency on the order of input scaffold assemblies. We remark that CAMSA can be utilized at different stages of the genome assembly process and be applied to assemblies of various genomic fragments, ranging from contigs to superscaffolds. In particular, CAMSA input can include results of other assembly reconciliation methods.

## 2. Methods

### Assembly Analysis and Visualization

For the purpose of comparative analysis and visualization of the input scaffold assemblies, CAMSA utilizes the *breakpoint graphs*, the data structure traditionally used for analysis of gene orders across multiple species [4]. We will refer to the breakpoint graph constructed on a set of scaffold assemblies as the *scaffold assembly graph* (SAG).

We start with the case of assemblies with no unoriented scaffolds. Then each assembly *A* can be viewed as a set of sequences of oriented scaffolds. We represent *A* as an individual *scaffold assembly graph* SAG(*A*) with two types of edges: directed edges (*scaffold edges*) encoding scaffolds in *A*, and undirected edges (*assembly edges*) representing scaffold adjacencies and connecting extremities (tails/heads) of the corresponding scaffold edges (Fig. 1A,B,C).

**Figure 1:**
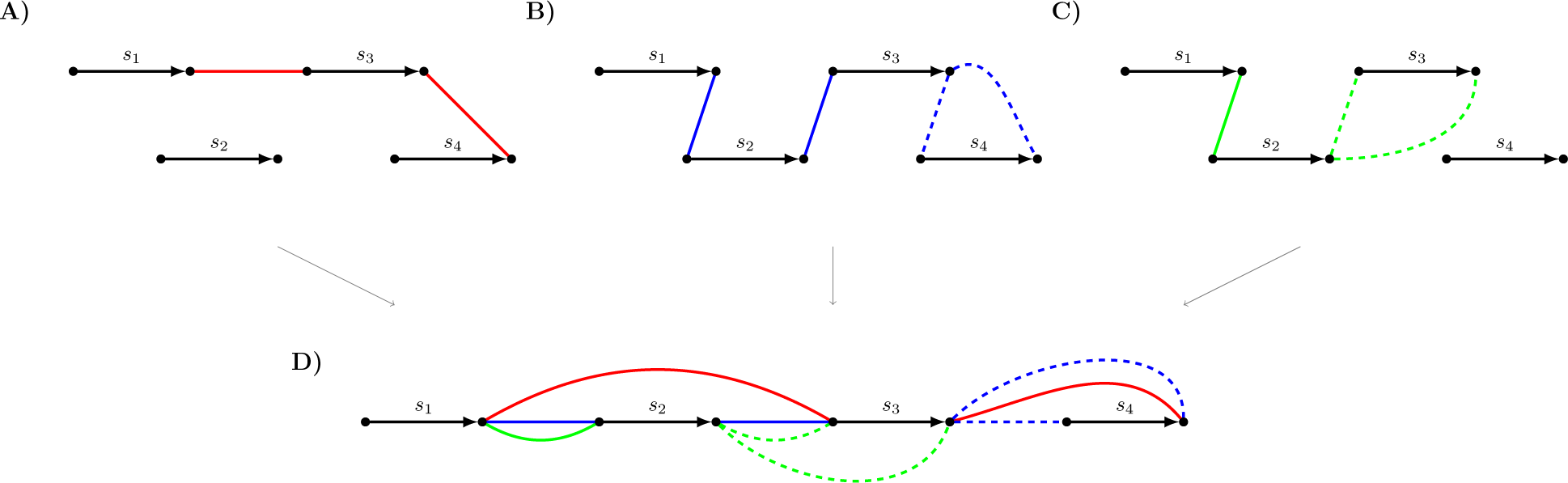
Individual scaffold assembly graphs for assemblies 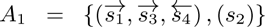 (red edges), 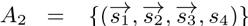 (blue edges), and 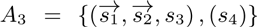 (green edges), and their scaffold assembly graph SAG(*A*_1_, *A*_2_, *A*_3_). scaffold edges are colored black. Actual assembly edges are shown as solid, while candidate assembly edges are shown as dashed. **A)** Individual scaffold assembly graph SAG(*A*_1_). **B)** Individual scaffold assembly graph SAG(*A*_2_). **C)** Individual scaffold assembly graph SAG(*A*_3_). **D)** scaffold assembly graph SAG(*A*_1_, *A*_2_, *A*_3_).

We find it convenient to refer to each assembly edge as an *assembly point*. Equivalently, an assembly point in *A* can be represented by an ordered pair of oriented scaffolds. We specify the orientation of a scaffold *s*, either by a sign (+*s* or *−s*) or by an overhead arrow 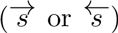. For example, 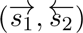 and 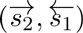 represent the same assembly point between scaffolds *s*_1_ and *s*_2_following each other head-to-head. Clearly, any assembly is completely defined by the set of its assembly points.

To construct the scaffold assembly graph SAG(*A*_1_,…, *A*_*k*_) of multiple input assemblies *A*_1_,…, *A*_*k*_, we represent them as individual graphs SAG(*A*_1_),…, SAG(*A*_*k*_), where the undirected edges in each SAG(*A*_*i*_) are colored into unique color. Then the graph SAG(*A*_1_,…, *A*_*k*_) can be viewed as the superposition of individual graphs SAG(*A*_1_),…, SAG(*A*_*k*_) and can be obtained by gluing the identically labeled directed edges. So the graph SAG(*A*_1_,…, *A*_*k*_) consists of (directed, labeled) scaffold edges encoding scaffolds and (undirected, unlabeled) assembly edges of *k* colors encoding assembly points in different input assemblies (Fig. 1D). We will refer to edges of color *A*_*i*_ (i.e., coming from SAG(*A*_*i*_)) as *A*_*i*_-edges. The assembly edges connecting the same two vertices *x* and *y* form the *multiedge {x, y}* in SAG(*A*_1_,…, *A*_*k*_). The *multicolor* of *{x, y}* is defined as the union of the colors of individual edges connecting *x* and *y*.

We define the (ordinary) *degree* odeg(*x*) of a vertex *x* in SAG(*A*_1_,…, *A*_*k*_) as the number of assembly edges incident to *x*. We distinguish it from the *multidegree* mdeg(*x*) defined as the number of adjacent vertices that are connected to *x* with assembly edges.

When all assemblies *A*_1_,…, *A*_*k*_ agree on a particular assembly point *{x, y}*, the graph SAG(*A*_1_,…, *A*_*k*_) contains a multi-edge *{x, y}* composed of edges of all *k* different colors. In other words, both vertices *x* and *y* in this case have degree *k* and multidegree 1. For a vertex *z* in SAG(*A*_1_,…, *A*_*k*_), odeg(*z*) ≠ *k* or mdeg(*z*) ≠ 1 indicate some type of inconsistency between the assemblies.

We classify an individual assembly points *{x, y}* as follows. Let *S* be the multicolor of the multiedge *{x, y}* in SAG(*A*_1_,…, *A*_*k*_).

- **unique** if *|S|* = 1, i.e., the assembly point *{x, y}* is present only in a single assembly;
- **in-conflicting** within assembly *A* ∈ *S* if *x* or *y* is incident any other *A*-edges besides *{x, y}* (e.g., Fig. 2C);
- **out-conflicting** if there exist two distinct assemblies *A* and *B* such that *A* contains *{x, y}* (i.e., *A ∈ S*), and *B* contains *{x, z}* with *z* ≠ *y* or *{y, z}* with *z* ≠ *x* (e.g., Fig. 2A).

**Figure 2:**
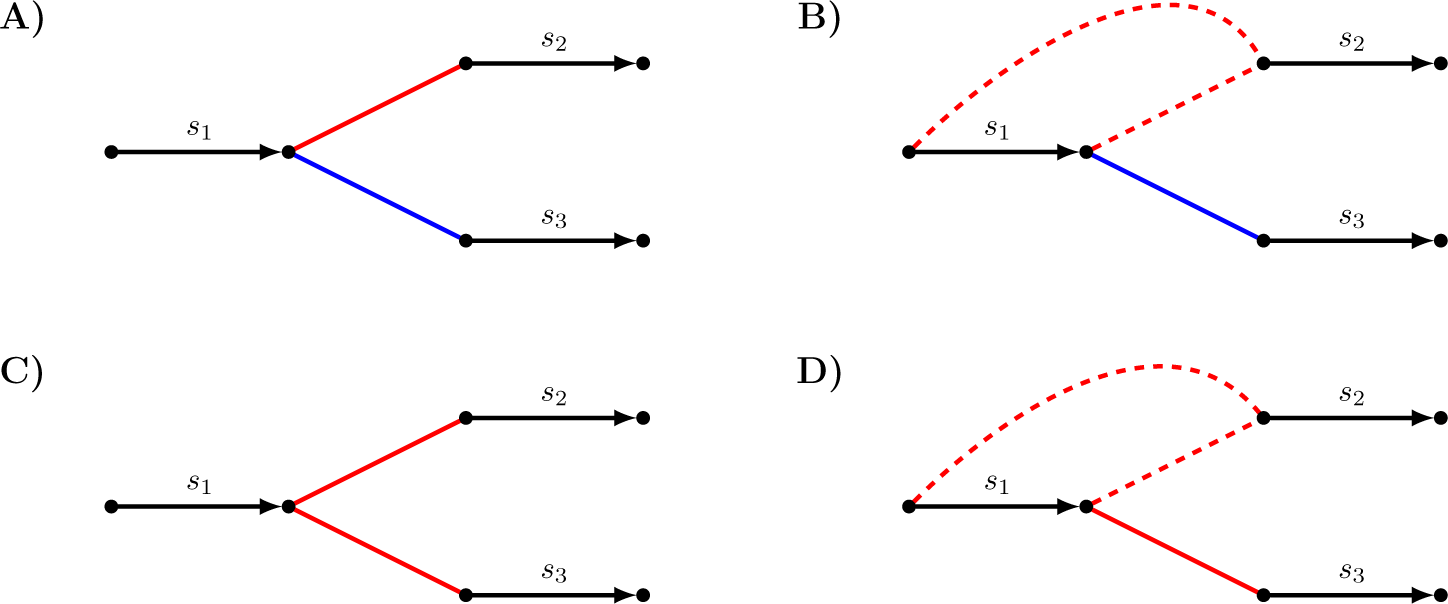
Illustration of various conflicts between assembly points of assemblies *A*_1_ (red edges) and *A*_2_ (blue edges).

### Dealing with Unoriented scaffolds

While conventional multiple breakpoint graphs are constructed for sequences of oriented genes, in CAMSA we extend scaffold assembly graphs to support assemblies that may include oriented as well as unoriented scaffolds.

In addition to (oriented) assembly points formed by pairs of oriented scaffolds, we now consider semi-oriented and unoriented assembly points.

A *semi-oriented* assembly point represents an adjacency between an oriented scaffold and an unoriented one. For example, 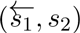 and 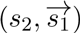 denote the same semi-oriented assembly point, where scaffold *s*_1_ is oriented and *s*_2_ is not (as emphasized by a missing overhead arrow). Similarly, an *unoriented* assembly point represents an adjacency between two unoriented scaffolds. For example, (*s*_1_, *s*_2_) and (*s*_2_, *s*_1_) denote the same unoriented assembly point between unoriented scaffolds *s*_1_ and *s*_2_.

We define a *realization* of an assembly point *p* as any oriented assembly point that can be obtained from *p* by orienting unoriented scaffolds. We denote the set of orientations of *p* as *R*(*p*). If *p* is oriented, then it has a single realization equal to *p* itself (i.e., *R*(*p*) = *{p}*); if *p* is semi-oriented, then it has two realizations (i.e., *|R*(*p*)*|* = 2); and if *p* is unoriented, then it has four realizations (i.e., *|R*(*p*)*|* = 4). For example,

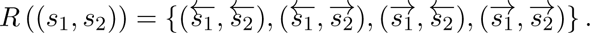

In the scaffold assembly graph, we add assembly edges encoding all realizations of semi-/un-oriented assembly points and refer to such edges as *candidate*, in contrast to *actual* assembly edges encoding oriented assembly points.

We extend the in-/out-conflicting classification to semi-oriented and unoriented assembly points as follows. An assembly point is *in-/out-conflicting* if all its realizations are such, except that we do not consider two realizations as in-conflicting to each other. Similarly, an assembly point is *in-/out-semiconflicting* if some but not all of its realizations are in-/out-conflicting (e.g., Fig. 2B,D illustrate pairs of out-semiconflicting and in-semiconflicting assembly points, respectively).

### Merging Assemblies

CAMSA can resolve conflicts in the input assemblies by merging them into a single (not self-confliciting) *merged assembly* that is most consistent with the input ones. The merged assembly is also used to determine orientation of (some) unoriented scaffolds in one input assemblies that is most confident and/or consistent with other input assemblies. In other words, the merged assembly helps to identify realizations of (some) semi-/un-oriented assembly points that are most consistent with other assemblies. Namely, for each semi-/un-oriented assembly point, the merged assembly contains either only one or none of its realizations; and in the former case, the included realization defines the most confident orientation of the corresponding unoriented scaffolds.

Assembly merging performed by CAMSA is based on how often each assembly point appears in the input assemblies as well as on the (optional) confidence of each such appearance. Namely, for each assembly point *p* in an input assembly *A*, CAMSA allows to specify the *confidence weight* cw_*A*_(*p*) from the interval [0, 1], which is then assigned to the corresponding assembly edge(s) (Fig. 3A). The confidence weights are expected to reflect the confidence level of the assembly methods in what they report as scaffold adjacencies (e.g., heuristic methods should probably have smaller confidence as compared to more reliable wet-lab techniques). By default, all actual assembly edges have the confidence weight equal 1, and all candidate assembly edges have weight 0.75 (these default values can be overwritten by the user).

**Figure 3:**
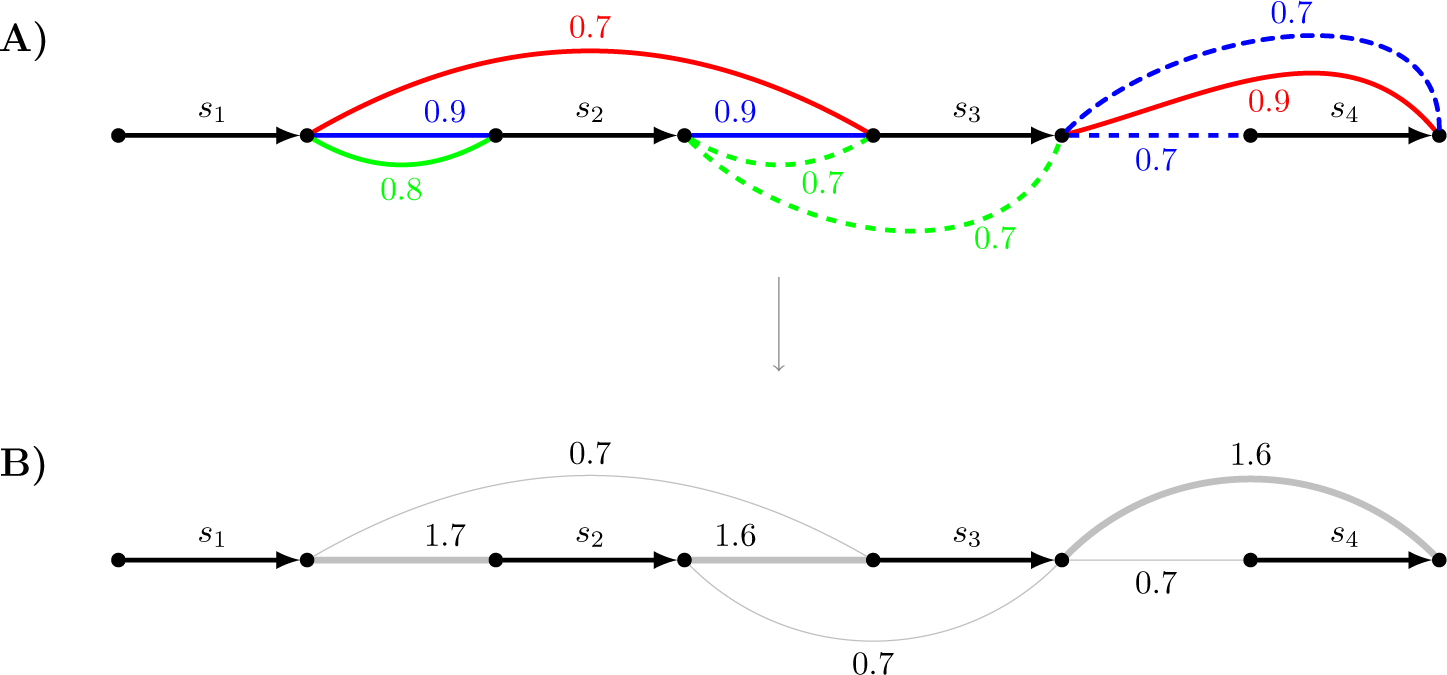
A) scaffold assembly graph SAG(*A*_1_, *A*_2_, *A*_3_), where assemblies *A*_1_, *A*_2_, and *A*_3_ are represented as red, blue, and green assembly edges, respectively, labeled with the corresponding confidence weights. **B)** Merged scaffold assembly graph MSAG(*A*_1_, *A*_2_, *A*_3_) obtained from SAG(*A*_1_, *A*_2_, *A*_3_) by replacing each assembly multi-edge with an ordinary edge of combined weight. The bold assembly edges represent the merged assembly computed by CAMSA.

For any oriented assembly *B* (viewed as a set of oriented assembly points), we define the *consistency score* cs_*B*_ (*A*) of an input assembly *A* with respect to *B* as 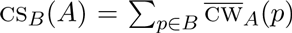,
where

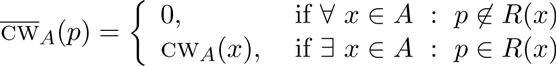

We pose the *assembly merging problem* (AMP) as follows.

**Problem 1** (Assembly Merging Problem, AMP). *Given assemblies A*_1_,…, *A*_*k*_ *of the same set of scaffolds S, find an assembly M of S containing only oriented assembly points such that*

i. *M is not self-conflicting (i.e., does not contain any in-conflicting assembly points);*
ii. 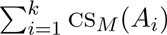 cs_*M*_ (*A*^*i*^) *is maximized;*
iii. *for every assembly point p* ∈ *A*_1_ *∪ · · · ∪ A*_*k*_, *at most one of its realizations is present in M* (*i.e.*, |*M* ∩ *R*(*p*)| ≤ 1).

For a solution *M* to the AMP, the condition (i) implies that the assembly edges in SAG(*M*) form a matching. Furthermore, *M* is assumed to correspond to the genome, which may be subject to additional constraints such as having all chromosomes linear (e.g., for vertebrate genomes) or having a single chromosome (e.g., for bacterial genomes). These constraints are translated for *M* as the absence in SAG(*M*) of cycles formed by alternating assembly and scaffold edges (for a unichromosomal circular genome, such a cycle can be present in SAG(*M*) only if it includes all scaffold edges).

To address the AMP, we start with construction of the (weighted) *merged scaffold assembly graph* MSAG(*A*_1_,…, *A*_*k*_) from SAG(*A*_1_,…, *A*_*k*_) by replacing each assembly multi-edge with an ordinary assembly edge of the weight equal the total weight of the corresponding multi-edge (Fig. 3). So, MSAG(*A*_1_,…, *A*_*k*_) is the graph with two types of edges: unweighted directed scaffolds edges and weighted undirected assembly edges. The AMP is then can be reformulated as the following *restricted maximum matching problem* (RMMP) on the graph *G* = MSAG(*A*_1_,…, *A*_*k*_):

**Problem 2** (Restricted Maximum Matching Problem, RMMP). *Given a merged scaffold assembly graph G, find a subset M of assembly edges in G such that*

i. *M is a matching;*
ii. *M has maximum weight;*
iii. *there are no cycles in SAG*(*M*).

Let *M* be a solution to the RMMP. Then the graph SAG(*M*) consists of scaffold edges forming a perfect matching and assembly edges from *M* forming a (possibly non-perfect) matching by the condition (i). Thus SAG(*M*) is formed by collection of paths and cycles, whose edges alternate between scaffold and assembly edges. Furthermore, by the condition (iii), SAG(*M*) consists entirely of alternating paths. A similar optimization problem, where the number of paths and the number cycles in the resulting SAG(*M*) are fixed, is known to be NP-complete [10], leaving a little hope for the RMMP to have a polynomial-time solution. Instead, CAMSA employs two merging heuristic solutions building upon the previously proposed algorithms [31, 10] as described below in this section.

**Greedy merging heuristics.** For a given merged scaffold assembly graph *G*, this strategy starts with the graph *H* consisting of scaffold edges from *G* and then iteratively enriches *H* with assembly edges so that no cycles are created in *H*. At any stage of this process, *H* is considered as a collection of alternating paths, some of which are merged into a longer path by adding a corresponding assembly edge. The paths to merge are selected based on the confidence weight of their linking assembly edge. The final graph *H* constructed this way defines *M* as the set of assembly edges in *H* (and so SAG(*M*) = *H*).

**Maximum matching heuristics.** For a given merged scaffold assembly graph *G*, this first computes the maximum weighted matching *M ′* formed by assembly edges of *G*. Namely, CAMSA employs the NetworkX library [19] implementation of the blossom algorithm [14] for computing *M ′*.^2^ For the maximum weighted matching *M ′*, CAMSA looks for cycles in SAG(*M ′*) (notice that all cycles in SAG(*M ′*) are vertex-disjoint) and removes an assembly edge of the lowest confidence weight from each such cycle. These edges are also removed from *M ′* to form *M* so that SAG(*M*) consists entirely of alternating paths.

We remark that before solving the RMMP for *G* = MSAG(*A*_1_,…, *A*_*k*_), CAMSA allows to remove assembly edges from *G* that have weight smaller than the *weight threshold* specified by the user (by default, this threshold is set to 0, i.e., no edges are removed). The removal of small-weighted assembly edges may be desirable if one wants to restrict attention only to assembly points of certain confidence level (e.g., assembly points coming either from individual highly-reliable assemblies or as a consensus from multiple assemblies). When such removal of low-confidence edges is performed, it is important to do so before (not after) solving the RMMP, since otherwise these edges may introduce a bias for an inclusion of high-confidence edges into the merged assembly *M*.

## 3. Structure of CAMSA Report

The results of comparative analysis and assembly merging performed by CAMSA are presented to the user in the form of an interactive *report*. The report is generated in a form of a JavaScript-powered HTML file, readily accessible for viewing/working in any modern Internet browser (for locally generated reports, Internet connection is not required). Many of the report sections are also available in the form of text files, making them accessible for machine processing. All tables in the report are powered by the DataTables JavaScript library [21], which provides flexible and dynamic filtering, sorting, and searching capabilities.

The first section of the CAMSA report presents aggregated characteristics of each input assemblies as compared to the others:

- the number of oriented, semi-oriented, and unoriented assembly points;
- the number of in-/out-conflicting assembly points;
- the number of in-/out-semiconflicting assembly points;
- the number of nonconflicting assembly points;
- the number of assembly points that participate in the merged assembly.

The second section of the CAMSA report focuses on consistency across various subsets of input assemblies. For each subset, it gives characteristics similar to the ones in the first section, but the values here are aggregated over all assemblies in the subset. The subsets are listed as a bar diagram in the descending order of the number of unique assembly points they contain (Fig. 4). Such statistics eliminates the need of running CAMSA separately on any assemblies subsets and allows the user to easily identify groups of assemblies that agree/conflict among themselves the most. We remark that each assembly point is counted only once: for the set of assemblies that contains this assembly point (but not for any its smaller subset). Since the the number of all subsets of input assemblies can be large, CAMSA allows the user to specify the number of top subsets to be displayed.

**Figure 4:**
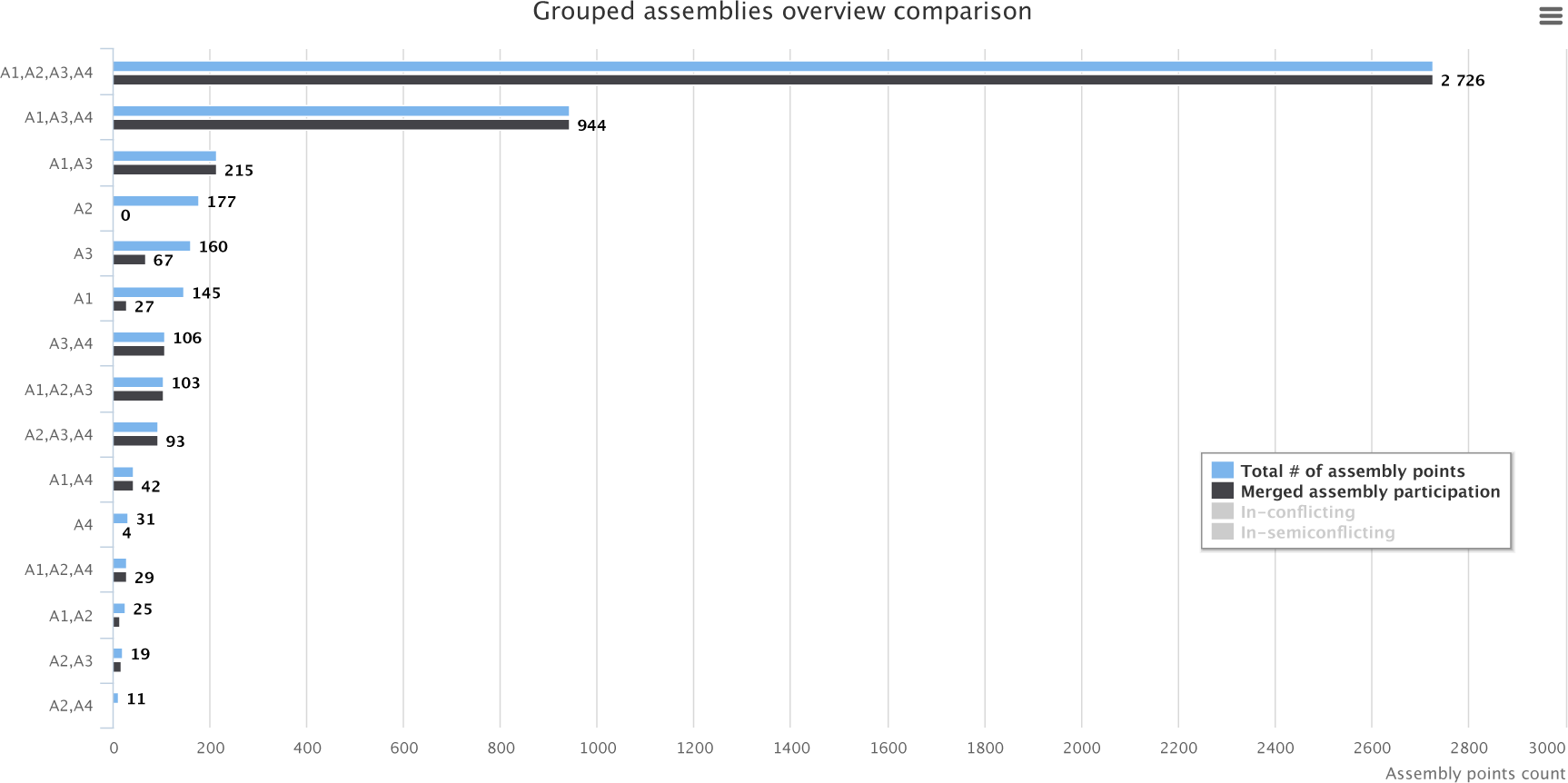
The second section of the CAMSA report for the scaffold assemblies of *H. sapiens Chr14* produced by ScaffMatch (*A*1), SGA (*A*2), SOAPdenovo2 (*A*3), and SSPACE (*A*4). For each subset of the assemblies *A*1, *A*2, *A*3, and *A***s**4**ca**,**ff**i**m**t**at**g**ch**iv[Ae1s] the number of assembly points that are unique to this subset; participate in the merged assembly; are inS-hcoownfl10 ictienntgrie;s and are in-semiconflicting.

The third section of the CAMSA report provides statistics for each assembly point within each assembly. Extensive interactive filtering allows the user to select assembly points of interest, as well as to export the filtered results, creating problem-/ region-/ fragment-focused analysis pipelines. We remark that statistical characteristics (e.g., whether an assembly point is in-/out-conflicting or in-/out-semiconflicting) are computed with respect to all of the input assemblies.

The fourth section of the CAMSA report provides statistics for each assembly point aggregated over all of the input assemblies (Fig. 5). In contrast to the third section, each assembly point is shown here exactly once, and the *sources* column shows the set of assemblies where this assembly point is present (e.g., in Fig. 5 the assembly point 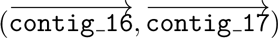 is present in *A*1, *A*2, and *A*3). Again, CAMSA provides extensive filtering to enable a focused analysis of assembly points of interest. The result of assembly points filtration can further be exported in the same format, which is utilized for CAMSA input files (i.e., list of assembly points in a tab-separated format).

**Figure 5:**
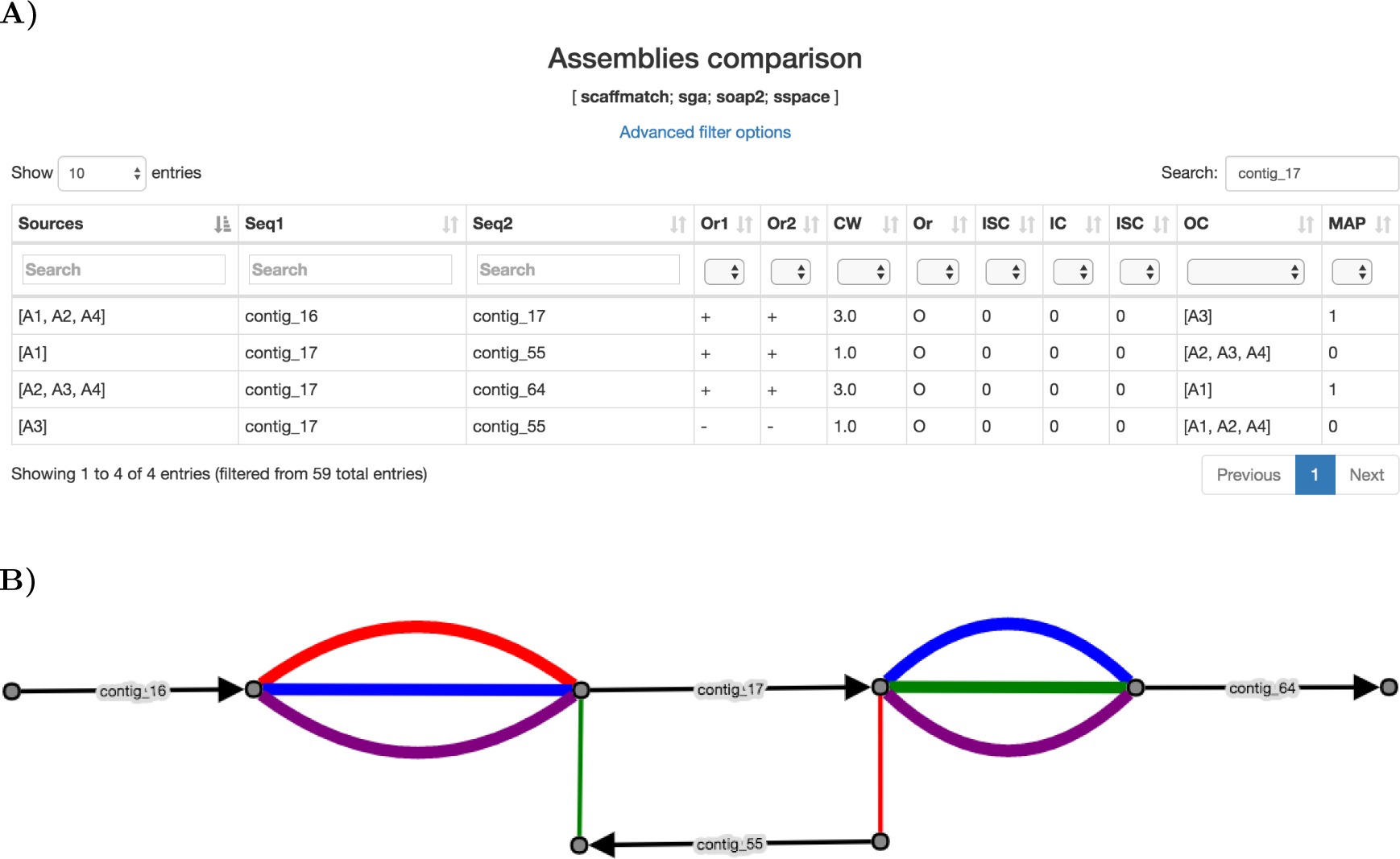
The fourth section of the CAMSA report for the scaffold assemblies of *S. aureus* produced by ScaffMatch (*A*1), SGA (*A*2), SOAPdenovo2 (*A*3), and SSPACE (*A*4). **A)** Table resulting from filtration and containing only assembly points involving scaffold contig 17. **B)** A subgraph of the scaffold assembly graph SAG(*A*1*, A*2*, A*3*, A*4) induced by the assembly points involving scaffold contig 17.

Besides the text-based representation and export, the CAMSA report also provides an interactive visualization and further graphical export of assembly points in the form of the scaffold assembly graph. A vector-based interactive graph visualization is created using the Cytoscape.js library [13]. This visualization has a dynamic graph layout and supports filtration of graph components. We allow the user to choose from several Cytoscape.js graph layouts; the default layout comes from [12].

At any point the current image of the scaffold assembly graph can be exported from the report into a PNG file.

The time required for graph visualization heavily depends on the chosen layout and the underlying graph complexity. In cases when visualization inside the report takes too much time, we provide the following workarounds. The assembly points can be exported in a text file and then converted into a DOT file describing the corresponding scaffold assembly graph, whose visualization then can be constructed with GraphViz [15]. Alternatively, one can choose to export the SAG subgraph induced by the filtered assembly points into a JSON file, which can further be processed with the desktop Cytoscape software [38].

## 4. Evaluation

While merging of multiple input scaffold assemblies is just one of the features of the CAMSA framework, it is the only one that resembles existing tools, namely those performing assembly reconciliation. We therefore feel obliged to compare its performance to such tools, even though we pose CAMSA as a meta-tool that can take as an input the results of various scaffolding methods, including assembly reconciliation tools.

We evaluated the assembly merging in CAMSA by running it on multiple scaffold assemblies of genomes of different sizes from the GAGE project [37]. While CAMSA can be used at any stage of genome scaffolding, in this evaluation we applied it to the results of initial scaffolding of contigs based on jumping libraries. We chose the following four scaffolders for performing such task: ScaffMatch [29], SOAPdenovo2 [27], SGA [39], and SSPACE [7]. The input to these scaffolders was formed by contigs and jumping libraries assembled and corrected by Allpaths-LG [16], which are provided by GAGE. The scaffold assemblies produced by these scaffolders were used as an input to CAMSA as well as to Metassembler [46] and GAM-NGS [44] assembly reconciliation tools.^3^ To demonstrate the advantages of CAMSA as a meta-tool, we also run it on the four aforementioned scaffold assemblies combined with the two reconciled assemblies produced by Metassembler and GAM-NGS, and denoted as CAMSA (+GM) in the evaluation results.

All tools were run on the same computer system with dual Intel Xeon E5-2670 2.6GHz 8-core processors and 64GB of RAM. First, we measured the running time of each tool. Then we assessed the quality of the resulting scaffold assemblies (formed by *merged scaffolds*) with the number of metrics computed by QUAST [18] with–scaffolds flag. Below we present most important metrics, while the complete QUAST reports for both input (Tables S8, S9, S10) and resulting scaffold assemblies (Tables S11, S12, S13) are provided in Supplement A. Namely, we mostly concern the following QUAST metrics:

- # contigs: in our evaluation, the contigs counted by QUAST correspond the merged scaffolds; so their number measures the contiguity of the resulting scaffold assemblies.
- # misassemblies (*miss.*): number of breakpoints in the merged scaffolds, for which the left and right flanking sequences align in the reference genome to different strands / chromosomes (inversions / translocations), or on the same stand and chromosome with a gap of *≥* 1000bp between each other (relocations).
- # local misassemblies (*local miss.*): number of relocations with a gap in the range from 85bp to 1000bp.
- NA50: the maximum length *L* such that the fragments of length *≥ L* obtained from the merged scaffolds by breaking them at misassembly sites cover at least 50% of assembly. NA75: similar to NA50, but with 75% coverage of the assembly.

Table 1 demonstrates that CAMSA is the fastest among the tools in comparison. We separately benchmarked the data preparation and processing. We remark that depending on the format of input scaffold assemblies as well as the overall assembly pipeline, the data preparation step may be not required or take significantly different time. For CAMSA in this evaluation data preparation involves the conversion of scaffold assemblies from FASTA format into the set of assembly points,^4^ using a utility script based on NUCmer software [23]. For GAM-NGS, one needs to align the jumping libraries onto the input scaffold assemblies as well as onto the intermediate reconciled assemblies (progressively generated from the input assemblies). The former alignments were treated as data preparation (since they may be readily available from the assembly pipeline), while the latter alignments are generally unavailable and thus were treated as data processing. For Metassembler, no data preparation was required since all alignments are performed internally.

**Table 1:** Running time of GAM-NGS, Metassembler, and CAMSA on scaffold assemblies produced by ScaffMatch, SOAPdenovo2, SGA, and SSPACE on three GAGE datasets. Time in parentheses is additional and corresponds to the data preparation. Best results are shown in bold.

|  | *S. aureus* | *R. sphaeroides* | *H. sapiens Chr14* |
| --- | --- | --- | --- |
| GAM-NGS | 4m25s (+2m3s) | 8m47s (+4m14s) | 1h29m (+43m) |
| Metassembler | 59m16s (+0s) | 1h48m53s (+0s) | 8h19m10s (+0s) |
| CAMSA | **2s (+3s)** | **2s (+10s)** | **48s (+59m)** |
| CAMSA (+GM) | **2s (+3s)** | **2s (+10s)** | 54s (+1h10m) |

Table 2 shows the quality of the scaffold assemblies produced by different tools. In all datasets, the assembly produced by CAMSA was either the best or very close to the best in each of the metrics. We remark that in some cases CAMSA (+GM) takes advantage of the reconciled assemblies and demonstrates better results than CAMSA. In other cases, however, having the reconciled assemblies turns out to be disadvantageous due to the elevated presence of misassemblies in them. This emphasizes the fact that assembly reconciliation/merging is sensitive to the quality of input assemblies and should be interpreted with caution. The comparative report in CAMSA can greatly help in identification of conflicting assembly points (indicating potential misassemblies), enabling their targeted analysis.

**Table 2:** Quality of the reconciled/merged scaffold assemblies constructed by GAM-NGS, Metassembler, and CAMSA from the scaffold assemblies produced by ScaffMatch, SOAPdenovo2, SGA, and SSPACE on three GAGE datasets. Best results are shown in bold.

| *S. aureus* |  |  |  |  |  |
| --- | --- | --- | --- | --- | --- |
|  | # contigs | # miss. | # local miss. | NA50 | NA75 |
| GAM-NGS | **6** | **0** | 6 | 1082860 | 1082860 |
| Metassembler | **6** | **0** | 3 | 1083010 | 1083010 |
| CAMSA | **6** | **0** | 3 | **1083448** | **1083448** |
| CAMSA (+GM) | **6** | **0** | **2** | 1083436 | 1083436 |

| *R. sphaeroides* |  |  |  |  |  |
| --- | --- | --- | --- | --- | --- |
|  | # contigs | # miss. | # local miss. | NA50 | NA75 |
| GAM-NGS | 16 | **6** | 17 | 3080645 | 3080645 |
| Metassembler | **9** | **6** | 17 | **3080845** | **3080845** |
| CAMSA | **9** | **6** | **9** | 2965313 | 2965313 |
| CAMSA (+GM) | 10 | **6** | **9** | 2964450 | 2964450 |

| *H. sapiens Chr14* |  |  |  |  |  |
| --- | --- | --- | --- | --- | --- |
|  | # contigs | # miss. | # local miss. | NA50 | NA75 |
| GAM-NGS | 128 | **83** | 543 | 2941846 | 1235019 |
| Metassembler | **93** | 94 | 528 | 2494911 | 1235460 |
| CAMSA | 109 | 91 | **485** | 2624904 | **1235471** |
| CAMSA (+GM) | 94 | 84 | 511 | **2979834** | **1235464** |

## 5. Discussion

CAMSA addresses the current deficiency of tools for automated comparison and analysis of multiple assemblies of the same set scaffolds. Since there exist numerous methods and techniques for scaffold assembly, identifying similarities and dissimilarities across assemblies produced by different methods is beneficial both for the developers of scaffold assembly algorithms and for the researchers focused on improving draft assemblies of specific organisms.

We remark that CAMSA expects as an input a list of assembly points, which differs from the output produced by some conventional scaffolding tools. This inspired us to develop a set of utility scripts that automate the input/output conversion process for CAMSA (e.g., from/to formats like FASTA, AGPv2, or GRIMM), and include them in the CAMSA distribution.

We further plan to enrich the graph-based analysis in CAMSA with various pattern matching techniques, enabling a better classification of assembly conflicts based on their origin (e.g., conflicting scaffold orders, wrong orientation of scaffolds, or different resolution of assemblies). We also plan on adding a *reference* mode, so that classification of assembly points in the input assemblies can be done with respect to a known reference genome, rather than just with respect to each other.

We also remark that CAMSA is currently utilized in the study of Anopheles mosquito genomes [33], where multiple research laboratories (including ours) work on improving the existing assemblies for a set of mosquito species.

## Footnotes

1 We remark that contigs can be viewed as scaffolds with no gaps. So, under scaffolds we understand both contigs and scaffolds.

2 The blossom algorithm computes a maximal weighted matching in a graph in *O*(*V*^3^) time, where *V* is the number of vertices.

3 We also considered GARM [30], but were unable to run it on any GAGE dataset, facing issues similar to those reported in [46].

4 We remark that conversion, for example, from NCBI AGPv2 format (rather than FASTA) would be much faster.

## Supplement A Evaluation details

### Datasets

Each dataset in the evaluation comes from the GAGE project and consists of the following files (the paths are relative to the root directory http://gage.cbcb.umd.edu/data/<genome>/, where <genome> is Staphylococcus aureus, Rhodobacter sphaeroides, and Hg chr14, respectively):

Reference genomic sequence file genome.fasta from Data.original/ directory;

Allpaths-LG corrected jumping library file shortjump {1,2}.fastq (and file longjump {1,2}.fastq, when available) from Data.allpathsCor.tgz archive;

Allpaths-LG assembled contigs from Assembly.tgz archive.

### Software

Data preparation, processing, and analysis in the evaluation were performed with the following software tools (particular versions are specified in parentheses):

1. Allpaths-LG (r52488) [16] (http://software.broadinstitute.org/allpaths-lg/blog/)
2. QUAST (v4.1) [18] (http://quast.sourceforge.net/)
3. Bowtie2 (2.2.9) [25] (http://bowtie-bio.sourceforge.net/bowtie2/)
4. Picard (1.129) (http://broadinstitute.github.io/picard)
5. Samtools (1.2) [26] (http://samtools.sourceforge.net/)
6. SGA (0.10.13) [39] (https://github.com/jts/sga)
7. SOAPdenovo2 (2.04-r240) [27] (https://github.com/aquaskyline/SOAPdenovo2)
8. ScaffMatch (0.9) [29] (http://alan.cs.gsu.edu/NGS/?q=content/scaffmatch)
9. Abyss (1.5.2) [40] (https://github.com/bcgsc/abyss)
10. SSPACE (3.0) [7] (http://www.baseclear.com/genomics/bioinformatics/basetools/SSPACE)
11. Metassembler (1.5) [46] (https://sourceforge.net/projects/metassembler/)
12. GAM-NGS (v1.1b) [44] (https://github.com/vice87/gam-ngs)
13. GARM (0.7.5) [30] (http://garm-meta-assem.sourceforge.net/)
14. CAMSA (1.0.0) (https://cblab.org/camsa)
15. NUCmer (3.1) [23] (http://mummer.sourceforge.net/)

### Experiments outline

For each GAGE dataset, the process of preparation, scaffold assembly, merging of the resulting scaffold assemblies, and their further analysis is outlined below:

1. Using QUAST, compute statistics of the Allpaths-LG contigs (Supplementary Tables S5, S6, S7).
2. Using Bowtie2, align the Allpaths-LG corrected shortjump (and longjump, when available) jumping libraries to the reference genome sequence.
3. Using Samtools and Picard tools, from the obtained reads-to-reference alignment determine jumping library orientation, the median insert size and its standard deviation (Supplementary Table S3).
4. (when required) Using Bowtie2, align the same corrected jumping libraries to the Allpaths-LG contigs (Supplementary Table S4 describes the alignment parameters).
5. scaffold the Allpaths-LG contigs with different scaffolders, using shortjump (and longjump, when available) jumping libraries.
6. Using QUAST, compute statistics of the obtained scaffold assemblies (Supplementary Tables S8, S9, S10).
7. Merge the obtained scaffold assemblies, using CAMSA, Metassembler, and GAM-NGS.
8. Using QUAST, compute statistics of the obtained merged scaffold assemblies (Supplementary Tables S11, S12, S13).

**Table S3:** Metrics computed with Picard’s CollectInsertSizeMetrics tool for jumping libraries in the three observed GAGE datasets.

| Dataset | Library | Orientation | Median insert size | Standard deviation | Size |
| --- | --- | --- | --- | --- | --- |
| *S. aureus* | shortjump | RF | 3609 | 265 | 475408 |
| *R. sphaeroides* | shortjump | RF | 3761 | 716 | 516804 |
|  |  | FR | 344 | 97 | 34296 |
| *H. sapiens Chr14* | shortjump | RF | 2702 | 256 | 2095666 |
|  |  | FR | 242 | 90 | 706331 |
|  | longjump | FR | 34755 | 7892 | 80331 |

**Table S4:** Bowtie2 parameters used in alignment jumping libraries to Allpaths-LG assembled contigs in the three observed GAGE dataset.

| Dataset | Library | Bowtie2 parameters |
| --- | --- | --- |
| *S. aureus* | shortjump | --minins 3300 --maxins 3900 --rf |
| *R. sphaeroides* | shortjump | --minins 3000 --maxins 4450 --rf |
| *H. sapience Chr14* | shortjump | --minins 2400 --maxins 3000 --rf |
|  | longjump | --minins 27000 --maxins 42500 |

**Table S5:** QUAST report for Allpaths-LG assembled contigs of *S. aureus*

| Assembly | *S. aureus* |
| --- | --- |
| # contigs ($\geq 1000$ bp) | 58 |
| # contigs ($\geq 5000$ bp) | 45 |
| # contigs ($\geq 10000$ bp) | 40 |
| # contigs ($\geq 25000$ bp) | 35 |
| # contigs ($\geq 50000$ bp) | 19 |
| Total length ($\geq 0$ bp) | 2870776 |
| Total length ($\geq 1000$ bp) | 2868733 |
| Total length ($\geq 5000$ bp) | 2840918 |
| Total length ($\geq 10000$ bp) | 2800923 |
| Total length ($\geq 25000$ bp) | 2715192 |
| Total length ($\geq 50000$ bp) | 2129621 |
| # contigs | 59 |
| Largest contig | 234488 |
| Total length | 2869581 |
| Reference length | 2903081 |
| GC (%) | 32.65 |
| Reference GC (%) | 32.73 |
| N50 | 96740 |
| NG50 | 96740 |
| N75 | 48304 |
| NG75 | 48304 |
| L50 | 10 |
| LG50 | 10 |
| L75 | 20 |
| LG75 | 20 |
| # misassemblies | 0 |
| # misassembled contigs | 0 |
| Misassembled contigs length | 0 |
| # local misassemblies | 0 |
| # unaligned contigs | 0 + 0 part |
| Unaligned length | 0 |
| Genome fraction (%) | 98.818 |
| Duplication ratio | 1.000 |
| # N's per 100 kbp | 1.50 |
| # mismatches per 100 kbp | 1.92 |
| # indels per 100 kbp | 1.01 |
| Largest alignment | 234488 |
| NA50 | 96740 |
| NGA50 | 96740 |
| NA75 | 45922 |
| NGA75 | 45922 |
| LA50 | 10 |
| LGA50 | 10 |
| LA75 | 21 |
| LGA75 | 21 |

**Table S6:**
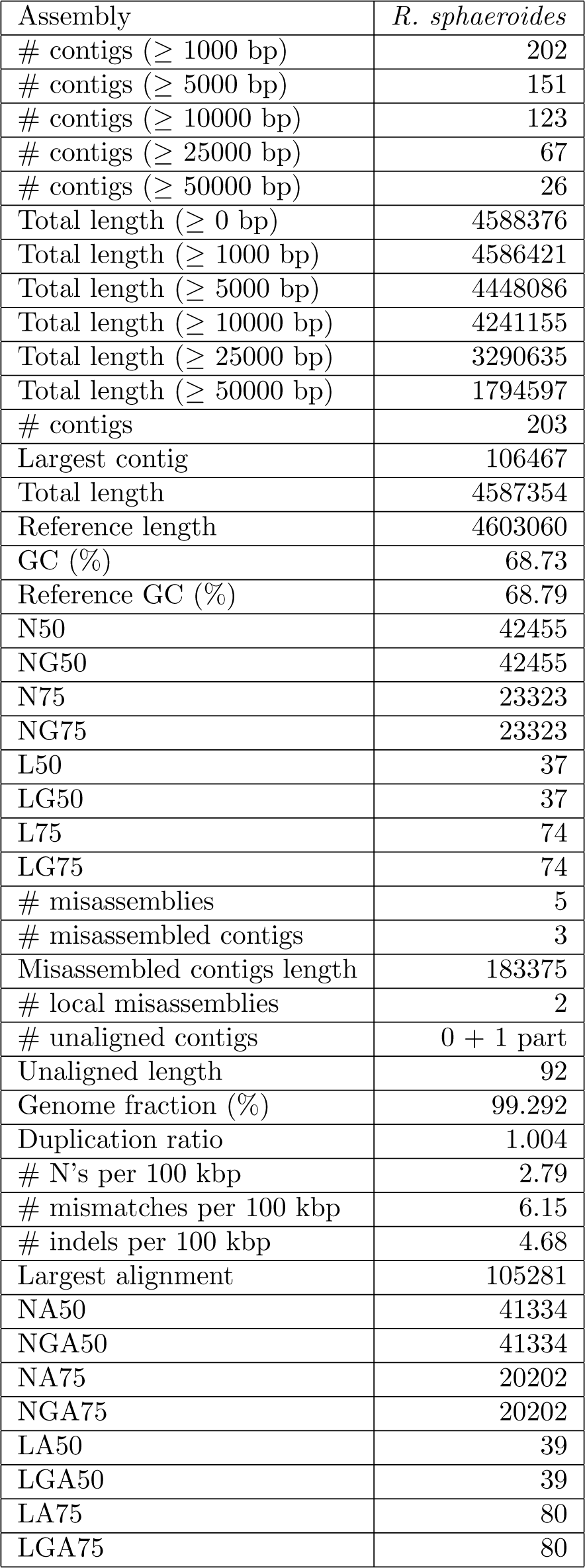
QUAST report for Allpaths-LG assembled contigs of *R. sphaeroides*

**Table S7:** QUAST report for Allpaths-LG assembled contigs of *H. sapiens Chr14*

| Assembly | *H. Sapiens Chr14* |
| --- | --- |
| # contigs ($\geq 1000$ bp) | 4383 |
| # contigs ($\geq 5000$ bp) | 2965 |
| # contigs ($\geq 10000$ bp) | 2256 |
| # contigs ($\geq 25000$ bp) | 1152 |
| # contigs ($\geq 50000$ bp) | 407 |
| Total length ($\geq 0$ bp) | 84461065 |
| Total length ($\geq 1000$ bp) | 84346908 |
| Total length ($\geq 5000$ bp) | 80952335 |
| Total length ($\geq 10000$ bp) | 75700427 |
| Total length ($\geq 25000$ bp) | 57826694 |
| Total length ($\geq 50000$ bp) | 31727320 |
| # contigs | 4469 |
| Largest contig | 240773 |
| Total length | 84416102 |
| Reference length | 107349540 |
| GC (%) | 40.77 |
| Reference GC (%) | 40.89 |
| N50 | 38359 |
| NG50 | 27960 |
| N75 | 20286 |
| NG75 | 5549 |
| L50 | 646 |
| LG50 | 995 |
| L75 | 1396 |
| LG75 | 2882 |
| # misassemblies | 51 |
| # misassembled contigs | 51 |
| Misassembled contigs length | 594366 |
| # local misassemblies | 138 |
| # unaligned contigs | 0 + 78 part |
| Unaligned length | 10377 |
| Genome fraction (%) | 78.466 |
| Duplication ratio | 1.002 |
| # N's per 100 kbp | 54.60 |
| # mismatches per 100 kbp | 67.25 |
| # indels per 100 kbp | 21.79 |
| Largest alignment | 240773 |
| NA50 | 38186 |
| NGA50 | 27586 |
| NA75 | 20019 |
| NGA75 | 5273 |
| LA50 | 647 |
| LGA50 | 999 |
| LA75 | 1404 |
| LGA75 | 2917 |

**Table S8:** QUAST report for 4 obtained scaffold assemblies on the *S. aureus* dataset

| Assembly | SGA | SSPACE | SOAPdenovo2 | ScaffMatch |
| --- | --- | --- | --- | --- |
| # contigs ($\geq 1000$ bp) | 16 | 8 | **6** | 7 |
| # contigs ($\geq 5000$ bp) | 10 | **6** | **6** | **6** |
| # contigs ($\geq 10000$ bp) | 7 | **6** | **6** | **6** |
| # contigs ($\geq 25000$ bp) | 7 | 6 | **5** | **5** |
| # contigs ($\geq 50000$ bp) | 6 | **4** | **4** | **4** |
| Total length ($\geq 0$ bp) | 2879803 | **2888431** | 2885823 | 2885463 |
| Total length ($\geq 1000$ bp) | 2879515 | **2887236** | 2884916 | 2884556 |
| Total length ($\geq 5000$ bp) | 2867655 | 2884858 | **2884916** | 2883180 |
| Total length ($\geq 10000$ bp) | 2847154 | 2884858 | **2884916** | 2883180 |
| Total length ($\geq 25000$ bp) | 2847154 | **2884858** | 2860116 | 2859903 |
| Total length ($\geq 50000$ bp) | 2808847 | 2820425 | **2821809** | 2821596 |
| # contigs | 16 | 8 | **6** | 7 |
| Largest contig | 1436473 | **1437245** | 1436067 | 1436757 |
| Total length | 2879515 | **2887236** | 2884916 | 2884556 |
| Reference length | 2903081 | 2903081 | 2903081 | 2903081 |
| GC (%) | 32.65 | 32.65 | 32.65 | 32.65 |
| Reference GC (%) | 32.73 | 32.73 | 32.73 | 32.73 |
| N50 | 690085 | 1093113 | **1096127** | 1094991 |
| NG50 | 690085 | 1093113 | **1096127** | 1094991 |
| N75 | 231306 | 1093113 | **1096127** | 1094991 |
| NG75 | 231306 | 1093113 | **1096127** | 1094991 |
| L50 | **2** | **2** | **2** | **2** |
| LG50 | **2** | **2** | **2** | **2** |
| L75 | 3 | **2** | **2** | **2** |
| LG75 | 3 | **2** | **2** | **2** |
| # misassemblies | **0** | **0** | **0** | **0** |
| # misassembled contigs | **0** | **0** | **0** | **0** |
| Misassembled contigs length | **0** | **0** | **0** | **0** |
| # local misassemblies | 1 | 2 | 6 | 3 |
| # scaffold gap size misassemblies | **23** | 41 | 28 | 31 |
| # unaligned contigs | **0 + 0 part** | **0 + 0 part** | **0 + 0 part** | **0 + 0 part** |
| Unaligned length | **0** | **0** | **0** | **0** |
| Genome fraction (%) | 98.832 | 98.824 | 98.817 | **98.847** |
| Duplication ratio | **1.004** | 1.006 | 1.006 | 1.005 |
| # N's per 100 kbp | **368.29** | 616.26 | 568.37 | 510.75 |
| # mismatches per 100 kbp | 3.83 | 3.97 | **3.17** | 4.46 |
| # indels per 100 kbp | 1.92 | **1.85** | 2.41 | 2.93 |
| Largest alignment | **1394578** | 1394477 | 1373048 | 1393974 |
| NA50 | 685027 | 1080969 | 1082860 | **1083369** |
| NGA50 | 685027 | 1080969 | 1082860 | **1083369** |
| NA75 | 231127 | 1080969 | 1082860 | **1083369** |
| NGA75 | 231127 | 1080969 | 1082860 | **1083369** |
| LA50 | **2** | **2** | **2** | **2** |
| LGA50 | **2** | **2** | **2** | **2** |
| LA75 | 3 | **2** | **2** | **2** |
| LGA75 | 3 | **2** | **2** | **2** |

**Table S9:** QUAST report for 4 obtained scaffold assemblies on the *R. sphaeroides* dataset

| Assembly | SGA | SSPACE | SOAPdenovo2 | ScaffMatch |
| --- | --- | --- | --- | --- |
| # contigs ($\geq 1000$ bp) | 79 | **15** | 16 | 16 |
| # contigs ($\geq 5000$ bp) | 55 | 9 | **8** | 9 |
| # contigs ($\geq 10000$ bp) | 50 | 9 | **8** | **8** |
| # contigs ($\geq 25000$ bp) | 39 | 8 | **7** | **7** |
| # contigs ($\geq 50000$ bp) | 23 | 7 | **6** | **6** |
| Total length ($\geq 0$ bp) | 4600980 | 4609201 | 4599261 | **4619707** |
| Total length ($\geq 1000$ bp) | 4600506 | 4607246 | 4598239 | **4619159** |
| Total length ($\geq 5000$ bp) | 4551065 | 4598883 | 4586030 | **4607565** |
| Total length ($\geq 10000$ bp) | 4515666 | **4598883** | 4586030 | 4598709 |
| Total length ($\geq 25000$ bp) | 4350981 | 4584344 | 4571440 | **4586110** |
| Total length ($\geq 50000$ bp) | 3784532 | 4550576 | 4537672 | **4552342** |
| # contigs | 79 | **16** | **16** | **16** |
| Largest contig | 471741 | 1605102 | **3186031** | 2544306 |
| Total length | 4600506 | 4608179 | 4598239 | **4619159** |
| Reference length | 4603060 | 4603060 | 4603060 | 4603060 |
| GC (%) | 68.72 | 68.73 | 68.72 | 68.73 |
| Reference GC (%) | 68.79 | 68.79 | 68.79 | 68.79 |
| N50 | 211952 | 1581585 | **3186031** | 2544306 |
| NG50 | 211952 | 1581585 | **3186031** | 2544306 |
| N75 | 73878 | **912982** | 911802 | 654158 |
| NG75 | 73878 | 912982 | 911802 | **914068** |
| L50 | 8 | 2 | **1** | **1** |
| LG50 | 8 | 2 | **1** | **1** |
| L75 | 18 | 3 | **2** | 3 |
| LG75 | 18 | 3 | **2** | **2** |
| # misassemblies | **5** | 6 | 6 | 6 |
| # misassembled contigs | 3 | 3 | 3 | **2** |
| Misassembled contigs length | **183375** | 335048 | 324030 | 323930 |
| # local misassemblies | **3** | 11 | 17 | 8 |
| # scaffold gap size misassemblies | **36** | 45 | 44 | 56 |
| # unaligned contigs | **0 + 0 part** | **0 + 0 part** | **0 + 0 part** | 0 + 1 part |
| Unaligned length | **0** | **0** | **0** | 4634 |
| Genome fraction (%) | 99.318 | 99.338 | 99.305 | **99.342** |
| Duplication ratio | **1.006** | 1.008 | **1.006** | 1.009 |
| # N's per 100 kbp | **336.16** | 470.88 | 338.39 | 681.05 |
| # mismatches per 100 kbp | 6.41 | 5.88 | 7.79 | **4.90** |
| # indels per 100 kbp | **6.67** | 7.35 | 8.16 | 7.22 |
| Largest alignment | 450352 | 1522122 | **3080645** | 2356266 |
| NA50 | 186972 | 1477526 | **3080645** | 2356266 |
| NGA50 | 186972 | 1477526 | **3080645** | 2356266 |
| NA75 | 58483 | **907420** | 886493 | 609106 |
| NGA75 | 58483 | **907420** | 886493 | 609106 |
| LA50 | 9 | 2 | **1** | **1** |
| LGA50 | 9 | 2 | **1** | **1** |
| LA75 | 21 | 3 | **2** | 3 |
| LGA75 | 21 | 3 | **2** | 3 |

**Table S10:** QUAST report for 4 obtained scaffold assemblies on the *H. sapiens Chr14* dataset

| Assembly | SSPACE | SOAPdenovo2 | ScaffMatch | SGA |
| --- | --- | --- | --- | --- |
| # contigs ($\geq 1000$ bp) | 469 | **116** | 231 | 1238 |
| # contigs ($\geq 5000$ bp) | 358 | **50** | 80 | 759 |
| # contigs ($\geq 10000$ bp) | 331 | **33** | 51 | 558 |
| # contigs ($\geq 25000$ bp) | 288 | **16** | 36 | 339 |
| # contigs ($\geq 50000$ bp) | 246 | **14** | 33 | 283 |
| Total length ($\geq 0$ bp) | 86098535 | 87702314 | 87845451 | **88055393** |
| Total length ($\geq 1000$ bp) | 86032471 | 87659994 | 87786387 | **87998362** |
| Total length ($\geq 5000$ bp) | 85823605 | **87532714** | 87507509 | 86938901 |
| Total length ($\geq 10000$ bp) | 85623892 | **87413294** | 87301150 | 85463731 |
| Total length ($\geq 25000$ bp) | 84949206 | **87150404** | 87074047 | 82203105 |
| Total length ($\geq 50000$ bp) | 83421289 | **87082426** | 86957082 | 80276434 |
| # contigs | 505 | **131** | 257 | 1288 |
| Largest contig | 3247728 | **27431299** | 11818947 | 1527937 |
| Total length | 86060510 | 87671707 | 87807154 | **88037622** |
| Reference length | 107349540 | 107349540 | 107349540 | 107349540 |
| GC (%) | 40.77 | 40.78 | 40.77 | 40.77 |
| Reference GC (%) | 40.89 | 40.89 | 40.89 | 40.89 |
| N50 | 479683 | **16717346** | 6850065 | 388109 |
| NG50 | 383690 | **12499179** | 4587587 | 296840 |
| N75 | 267450 | **9680384** | 2625940 | 178997 |
| NG75 | 92980 | **2923951** | 1342148 | 48481 |
| L50 | 50 | **2** | 5 | 67 |
| LG50 | 75 | **3** | 7 | 95 |
| L75 | 108 | **4** | 10 | 148 |
| LG75 | 204 | **7** | 17 | 288 |
| # misassemblies | **63** | 82 | 135 | 73 |
| # misassembled contigs | 42 | **18** | 26 | 60 |
| Misassembled contigs length | 20029844 | 83944488 | 80703080 | **15643725** |
| # local misassemblies | **460** | 540 | 543 | 292 |
| # scaffold gap size misassemblies | 2647 | 3649 | 2680 | **2200** |
| # unaligned contigs | 0 + 4 part | **0 + 3 part** | 0 + 4 part | 0 + 24 part |
| Unaligned length | **2008** | 3964 | 11617 | 5680 |
| Genome fraction (%) | 78.492 | 78.387 | **78.496** | 78.459 |
| Duplication ratio | **1.021** | 1.042 | 1.042 | 1.045 |
| # N's per 100 kbp | **1960.83** | 3823.84 | 3906.84 | 4188.37 |
| # mismatches per 100 kbp | 68.21 | 119.21 | **67.34** | 68.82 |
| # indels per 100 kbp | 23.21 | **22.00** | 23.25 | 22.94 |
| Largest alignment | 1734949 | **6288691** | 3518880 | 1493355 |
| NA50 | 414636 | **2941846** | 1399071 | 326554 |
| NGA50 | 333185 | **1965866** | 1001085 | 227608 |
| NA75 | 215879 | **1418204** | 554190 | 126637 |
| NGA75 | 59495 | **346413** | 155068 | 13585 |
| LA50 | 58 | **11** | 22 | 76 |
| LGA50 | 87 | **16** | 31 | 111 |
| LA75 | 127 | **23** | 48 | 184 |
| LGA75 | 251 | **46** | 96 | 497 |

**Table S11:** QUAST report for 4 merged scaffold assemblies on the *S. aureus* dataset.

| Assembly | Metassembler | GAM-NGS | CAMSA | CAMSA (+GM) |
| --- | --- | --- | --- | --- |
| # contigs ($\geq 1000$ bp) | **6** | **6** | **6** | **6** |
| # contigs ($\geq 5000$ bp) | **6** | **6** | **6** | **6** |
| # contigs ($\geq 10000$ bp) | **6** | **6** | **6** | **6** |
| # contigs ($\geq 25000$ bp) | **5** | **5** | 6 | 6 |
| # contigs ($\geq 50000$ bp) | **4** | **4** | **4** | **4** |
| Total length ($\geq 0$ bp) | 2886309 | 2885823 | 2887925 | **2889036** |
| Total length ($\geq 1000$ bp) | 2886309 | 2884916 | 2886730 | **2888129** |
| Total length ($\geq 5000$ bp) | 2886309 | 2884916 | 2886730 | **2888129** |
| Total length ($\geq 10000$ bp) | 2886309 | 2884916 | 2886730 | **2888129** |
| Total length ($\geq 25000$ bp) | 2861509 | 2860116 | 2886730 | **2888129** |
| Total length ($\geq 50000$ bp) | 2823202 | 2821809 | 2822351 | **2823801** |
| # contigs | **6** | **6** | **6** | **6** |
| Largest contig | 1436492 | 1436067 | 1436593 | **1437308** |
| Total length | 2886309 | 2884916 | 2886730 | **2888129** |
| Reference length | 2903081 | 2903081 | 2903081 | 2903081 |
| GC (%) | 32.65 | 32.65 | 32.65 | 32.65 |
| Reference GC (%) | 32.73 | 32.73 | 32.73 | 32.73 |
| N50 | **1096942** | 1096127 | 1095842 | 1096621 |
| NG50 | **1096942** | 1096127 | 1095842 | 1096621 |
| N75 | **1096942** | 1096127 | 1095842 | 1096621 |
| NG75 | **1096942** | 1096127 | 1095842 | 1096621 |
| L50 | **2** | **2** | **2** | **2** |
| LG50 | **2** | **2** | **2** | **2** |
| L75 | **2** | **2** | **2** | **2** |
| LG75 | **2** | **2** | **2** | **2** |
| # misassemblies | **0** | **0** | **0** | **0** |
| # misassembled contigs | **0** | **0** | **0** | **0** |
| Misassembled contigs length | **0** | **0** | **0** | **0** |
| # local misassemblies | 3 | 6 | 3 | **2** |
| # scaffold gap size misassemblies | 31 | **28** | 35 | 36 |
| # unaligned contigs | **0 + 0 part** | **0 + 0 part** | **0 + 0 part** | **0 + 0 part** |
| Unaligned length | **0** | **0** | **0** | **0** |
| Genome fraction (%) | 98.822 | 98.817 | 98.833 | **98.847** |
| Duplication ratio | **1.006** | **1.006** | **1.006** | 1.007 |
| # N's per 100 kbp | 607.70 | **568.37** | 595.55 | 633.84 |
| # mismatches per 100 kbp | 4.15 | **3.17** | 3.66 | 4.53 |
| # indels per 100 kbp | 2.37 | 2.41 | **2.27** | 2.61 |
| Largest alignment | 1374235 | 1373048 | 1373608 | **1394118** |
| NA50 | 1083010 | 1082860 | **1083448** | 1083436 |
| NGA50 | 1083010 | 1082860 | **1083448** | 1083436 |
| NA75 | 1083010 | 1082860 | **1083448** | 1083436 |
| NGA75 | 1083010 | 1082860 | **1083448** | 1083436 |
| LA50 | **2** | **2** | **2** | **2** |
| LGA50 | **2** | **2** | **2** | **2** |
| LA75 | **2** | **2** | **2** | **2** |
| LGA75 | **2** | **2** | **2** | **2** |

**Table S12:** QUAST report for 4 merged scaffold assemblies on the *R. sphaeroides* dataset.

| Assembly | Metassembler | GAM-NGS | CAMSA | CAMSA (+GM) |
| --- | --- | --- | --- | --- |
| # contigs ($\geq 1000$ bp) | **9** | 16 | **9** | 10 |
| # contigs ($\geq 5000$ bp) | **8** | **8** | **8** | **8** |
| # contigs ($\geq 10000$ bp) | **7** | 8 | **7** | **7** |
| # contigs ($\geq 25000$ bp) | **6** | 7 | **6** | **6** |
| # contigs ($\geq 50000$ bp) | **5** | 6 | **5** | **5** |
| Total length ($\geq 0$ bp) | 4611278 | 4599261 | **4619613** | 4618974 |
| Total length ($\geq 1000$ bp) | 4610804 | 4598239 | **4618591** | 4617952 |
| Total length ($\geq 5000$ bp) | 4609674 | 4586030 | **4617443** | 4615674 |
| Total length ($\geq 10000$ bp) | 4600818 | 4586030 | **4608585** | 4606816 |
| Total length ($\geq 25000$ bp) | 4586228 | 4571440 | **4593951** | 4592182 |
| Total length ($\geq 50000$ bp) | 4552460 | 4537672 | **4560183** | 4558414 |
| # contigs | **9** | 16 | **9** | 10 |
| Largest contig | 3186910 | 3186031 | **3193211** | 3191922 |
| Total length | 4610804 | 4598239 | **4618591** | 4617952 |
| Reference length | 4603060 | 4603060 | 4603060 | 4603060 |
| GC (%) | 68.72 | 68.72 | 68.73 | 68.73 |
| Reference GC (%) | 68.79 | 68.79 | 68.79 | 68.79 |
| N50 | 3186910 | 3186031 | **3193211** | 3191922 |
| NG50 | 3186910 | 3186031 | **3193211** | 3191922 |
| N75 | 912794 | 911802 | **913444** | 913317 |
| NG75 | 912794 | 911802 | **913444** | 913317 |
| L50 | **1** | **1** | **1** | **1** |
| LG50 | **1** | **1** | **1** | **1** |
| L75 | **2** | **2** | **2** | **2** |
| LG75 | **2** | **2** | **2** | **2** |
| # misassemblies | **6** | **6** | **6** | **6** |
| # misassembled contigs | **2** | 3 | **2** | **2** |
| Misassembled contigs length | 336947 | **324030** | 337651 | 337303 |
| # local misassemblies | 17 | 17 | **9** | **9** |
| # scaffold gap size misassemblies | 47 | **44** | 58 | 58 |
| # unaligned contigs | 0 + 1 part | **0 + 0 part** | 0 + 1 part | 0 + 1 part |
| Unaligned length | 4634 | **0** | 4636 | 4636 |
| Genome fraction (%) | 99.317 | 99.305 | 99.335 | **99.337** |
| Duplication ratio | 1.008 | **1.006** | 1.009 | 1.009 |
| # N's per 100 kbp | 621.87 | **338.39** | 679.10 | 665.36 |
| # mismatches per 100 kbp | 7.77 | 7.77 | **4.42** | **4.42** |
| # indels per 100 kbp | 8.16 | 8.16 | **7.28** | 7.33 |
| Largest alignment | **3080845** | 3080645 | 2964686 | 2964450 |
| NA50 | **3080845** | 3080645 | 2964686 | 2964450 |
| NGA50 | **3080845** | 3080645 | 2964686 | 2964450 |
| NA75 | 886881 | 886493 | **907644** | 907562 |
| NGA75 | 886881 | 886493 | **907644** | 907562 |
| LA50 | **1** | **1** | **1** | **1** |
| LGA50 | **1** | **1** | **1** | **1** |
| LA75 | **2** | **2** | **2** | **2** |
| LGA75 | **2** | **2** | **2** | **2** |

**Table S13:** QUAST report for 4 merged scaffold assemblies on the *H. sapience Chr14* dataset.

| Assembly | Metassembler | GAM-NGS | CAMSA | CAMSA (+GM) |
| --- | --- | --- | --- | --- |
| # contigs ($\geq 1000$ bp) | 86 | 113 | 94 | **82** |
| # contigs ($\geq 5000$ bp) | 39 | 48 | 37 | **36** |
| # contigs ($\geq 10000$ bp) | 25 | 30 | 26 | **24** |
| # contigs ($\geq 25000$ bp) | 13 | 16 | 17 | **12** |
| # contigs ($\geq 50000$ bp) | 12 | 14 | 16 | **11** |
| Total length ($\geq 0$ bp) | 87389910 | 87446601 | 87600310 | **87616405** |
| Total length ($\geq 1000$ bp) | 87380181 | 87404281 | 87552837 | **87571603** |
| Total length ($\geq 5000$ bp) | 87288768 | 87279334 | 87445013 | **87477118** |
| Total length ($\geq 10000$ bp) | 87192623 | 87149702 | 87370005 | **87392787** |
| Total length ($\geq 25000$ bp) | 87012955 | 86929915 | **87228898** | 87210713 |
| Total length ($\geq 50000$ bp) | 86983336 | 86861937 | **87199315** | 87181114 |
| # contigs | **93** | 128 | 109 | 94 |
| Largest contig | 27384426 | **27431299** | 22482944 | 27424679 |
| Total length | 87385570 | 87415994 | 87565052 | **87581387** |
| Reference length | 107349540 | 107349540 | 107349540 | 107349540 |
| GC (%) | 40.78 | 40.79 | 40.77 | 40.77 |
| Reference GC (%) | 40.89 | 40.89 | 40.89 | 40.89 |
| N50 | 22403791 | 16717346 | 13553269 | **22473965** |
| NG50 | 12475515 | 12499179 | 10575908 | **15417218** |
| N75 | 9667993 | 9680384 | 9654501 | **9681009** |
| NG75 | 3752984 | 2142674 | 3315947 | **5601426** |
| L50 | **2** | **2** | **2** | **2** |
| LG50 | **3** | **3** | **3** | **3** |
| L75 | **4** | **4** | **4** | **4** |
| LG75 | 6 | 8 | 7 | **5** |
| # misassemblies | 94 | **83** | 91 | 84 |
| # misassembled contigs | **12** | 18 | 18 | 13 |
| Misassembled contigs length | 85384146 | 83686313 | **78882289** | 85586152 |
| # local misassemblies | 528 | 543 | **485** | 511 |
| # scaffold gap size misassemblies | **2964** | 3639 | 3225 | 3303 |
| # unaligned contigs | **0 + 2 part** | 0 + 3 part | 0 + 3 part | **0 + 2 part** |
| Unaligned length | **3716** | 3964 | 8500 | 5087 |
| Genome fraction (%) | 78.449 | 78.161 | **78.486** | 78.485 |
| Duplication ratio | **1.038** | 1.042 | 1.039 | 1.040 |
| # N's per 100 kbp | **3470.36** | 3820.26 | 3629.33 | 3655.38 |
| # mismatches per 100 kbp | 119.28 | 119.05 | 67.82 | **67.71** |
| # indels per 100 kbp | 23.19 | **22.01** | 22.54 | 22.39 |
| Largest alignment | 6291358 | 6288691 | 5567517 | **6299514** |
| NA50 | 2494911 | 2941846 | 2624904 | **2979834** |
| NGA50 | 1836234 | 1836096 | 2164762 | **2182939** |
| NA75 | 1235460 | 1235019 | **1235471** | 1235464 |
| NGA75 | 272697 | **337612** | 272875 | 327733 |
| LA50 | 12 | **11** | 13 | **11** |
| LGA50 | 17 | 16 | 17 | **14** |
| LA75 | 25 | 24 | 24 | **21** |
| LGA75 | 51 | 48 | 50 | **46** |

